# MetDeeCINE: Deciphering Metabolic Regulation through Deep Learning and Multi-Omics

**DOI:** 10.1101/2025.03.24.645125

**Authors:** Takumi Ito, Satoshi Ohno, Yiran Wang, Saori Uematsu, Shinya Kuroda, Hideyuki Shimizu

**Affiliations:** Department of AI Systems Medicine, M&D Data Science Center, Institute of Integrated Research, Institute of Science Tokyo, Tokyo, Japan; School of Medicine, Faculty of Medicine, Institute of Science Tokyo, Tokyo, Japan; Department of Computational Biology and Medical Sciences, Graduate School of Frontier Sciences, The University of Tokyo, Tokyo, Japan; Department of Biological Sciences, Graduate School of Science, The University of Tokyo, Tokyo, Japan; Graduate School of Medical and Dental Sciences, Institute of Science Tokyo, Tokyo, Japan

**Keywords:** Multi-omics analysis, Metabolic engineering, Machine learning, Metabolism-informed neural network, XAI (Explainable Artificial Intelligence)

## Abstract

Metabolism, the biochemical reaction network within cells, is crucial for life, health, and disease. Recent advances in multi-omics technologies, enabling the simultaneous measurement of transcripts, proteins, and metabolites, provide unprecedented opportunities to comprehensively analyze metabolic regulation. However, effectively integrating these diverse data types to decipher the complex interplay between enzymes and metabolites remains a significant challenge due to the extensive data requirements of kinetic modeling approaches and the limited interpretability of machine learning approaches. Here, we present MetDeeCINE, a novel explainable deep learning framework that predicts the quantitative relationship between each enzyme and metabolite from proteomic and metabolomic data. We demonstrate that our newly developed Metabolism-informed Graph Neural Network (MiGNN), a core component of MetDeeCINE that is guided by the stoichiometric information of metabolic reactions, outperforms other machine learning models in predicting concentration control coefficients (CCCs) using data obtained from kinetic models of *E. coli*. Notably, MetDeeCINE, even without explicit information on allosteric regulation, can identify key distant enzymes that predominantly control the steady-state concentrations of specific metabolites. Application of MetDeeCINE to mouse liver multi-omics experimental data further demonstrated its ability to generate biologically meaningful predictions through identifying a rate-limiting enzyme of gluconeogenesis associated with obesity, consistent with existing knowledge. MetDeeCINE offers a scalable and interpretable approach for deciphering complex metabolic regulation from multi-omics data, with broad applications in disease research, drug discovery, and metabolic engineering.

## Introduction

Intracellular metabolism is an essential process for sustaining life, synthesizing key building blocks of cells (e.g., membranes, proteins, and nucleic acids), and generating energy storage molecules such as ATP. Metabolic dysregulation has been implicated in a wide range of diseases, including enzyme deficiencies, obesity, type 2 diabetes, cancer, and cardiovascular diseases^1–3^. Recent evidence suggests that elevated levels of specific metabolites, exceeding their normal physiological concentrations, promote disease progression. For example, oncometabolites in cancer highlight the potential of metabolic abnormalities as therapeutic targets^4^. Furthermore, in the field of metabolic engineering, the design of metabolic pathways to maximize the production of targeted compounds is an active area of research^5^. Thus, metabolism plays a central role in biology, medicine, and engineering, and a deeper, quantitative understanding of its regulatory mechanisms is crucial for breakthroughs in these fields.

Metabolism is regulated by a complex network spanning multiple omics layers, such as mRNA (transcriptome), proteins (proteome), and metabolites (metabolome), which are tightly interconnected. Each metabolite is converted from a substrate to a product by an enzymatic reaction, and the reaction rate (metabolic flux) depends on the amount of enzyme and the concentration of substrates. Moreover, certain metabolites, such as ATP and NADH, not only act as cofactors that affect the reaction rate in many metabolic processes but also exert allosteric effects on reactions elsewhere in the metabolic network^6^. Therefore, metabolic regulation is nonlinear, and alterations in the abundance of a particular RNA or enzyme can potentially impact the entire metabolic network^7^. Thus, methods that integrate multi-omics data are essential to achieve a systems-level, quantitative understanding of metabolic regulation.

Traditionally, kinetic modeling approaches using mathematical models have been employed for quantitative analysis of metabolic regulation from multi-omics data. In this approach, the metabolic network is represented as a system of ordinary differential equations (ODEs). Through Metabolic Control Analysis (MCA)^8,9^, kinetic models can be used to compute flux control coefficients (FCCs) and concentration control coefficients (CCCs), which quantify how changes in enzyme activity affect individual reaction rates and metabolite concentrations. These coefficients enable the identification of rate-limiting enzymes for the production of target compounds or therapeutic targets in metabolic diseases^10,11^. However, a critical limitation of this kinetic modeling approach is the requirement for detailed information for each metabolic reaction and the difficulty in scaling up owing to the stability of ODEs^12^. This detailed information is often unavailable, particularly for less-characterized organisms or pathways, limiting the applicability of kinetic modeling. Derived methods, such as ORACLE^13^ and OMELET^14^, incorporate thermodynamic constraints or Bayesian theory to predict FCCs and CCCs or infer the influence of each enzyme or metabolite on flux, without explicitly constructing the ODE model. Nevertheless, these methods still require extensive prior information, including reaction fluxes, allosteric regulators, and standard free energy changes, which limits the scale of the data or metabolic pathways to which they can be applied. Consequently, recent multi-omics studies of metabolism on human and mouse data often result in merely comparing each omics-layer rather than performing integrative analyses^15–18^.

With recent advancements in machine learning, various machine learning methods that utilize multi-omics data for metabolism have been reported. For instance, linear regression, multilayer perceptrons (MLPs), and random forest approaches have been used to predict the metabolome from proteomic data^19^ or to predict metabolic fluxes from transcriptomic, proteomic, and metabolomic data^20,21^. Although these methods do not rely on prior knowledge of metabolic networks or kinetic parameters, it remains unclear whether these model predictions accurately reflect the underlying mechanisms of metabolic regulation. While it is possible to identify important features, for instance, by computing SHAP values^22^ or regression coefficients in linear models, inferring how those features exert their influence through specific metabolic pathways remains challenging. Moreover, the predictive accuracy of purely data-driven approaches is limited by the high dimensionality and relatively small sample sizes typical of metabolic multi-omics datasets. Determining whether a prediction is biologically relevant or overfitted to noise remains challenging. A new approach, integrating metabolic network structure with multi-omics data, is eagerly awaited to bridge the gap between the detailed, mechanistic insights of kinetic modeling and the scalability of data-driven machine learning, enabling a comprehensive understanding of metabolic regulation at the system level.

In this study, we present MetDeeCINE (Metabolic analytical framework using Deep learning for Comprehensive Inference of multi-omics NEtwork), a novel and universally applicable framework for quantitatively analyzing metabolic regulation networks from multi-omics data (Fig. 1). MetDeeCINE predicts concentration control coefficients (CCCs) matrix which represent how changes in enzyme activities in a steady-state condition affect metabolite concentrations, from multi-omics data (Methods). MetDeeCINE combines concepts from traditional approaches based on metabolic control analysis with machine learning to elucidate the quantitative relationships between enzymes and metabolites at an omics scale, using only measured data and readily available stoichiometric information about metabolic reactions. Unlike previous approaches, MetDeeCINE does not require detailed kinetic parameters or explicit information on allosteric regulation, making it applicable to a wide range of organisms and pathways. We demonstrate that our metabolism-informed Graph Neural Network (MiGNN), a specialized deep learning model that incorporates stoichiometric information, achieves superior predictive performance. Using both simulation and experimental data, MetDeeCINE accurately predicts how enzymes can influence metabolite concentrations across the entire network, regardless of the network distance. Because MetDeeCINE can be applied to any species or metabolic pathway, it promises to enable detailed elucidation of metabolic regulation mechanisms at an omics scale that was previously difficult to achieve through independent analyses of single omics-layers. MetDeeCINE has broad applications, including the discovery of therapeutic targets in metabolic diseases, uncovering unknown metabolic regulatory mechanisms in biochemistry, or optimizing microbial engineering for improved compound production.

**Figure 1.**
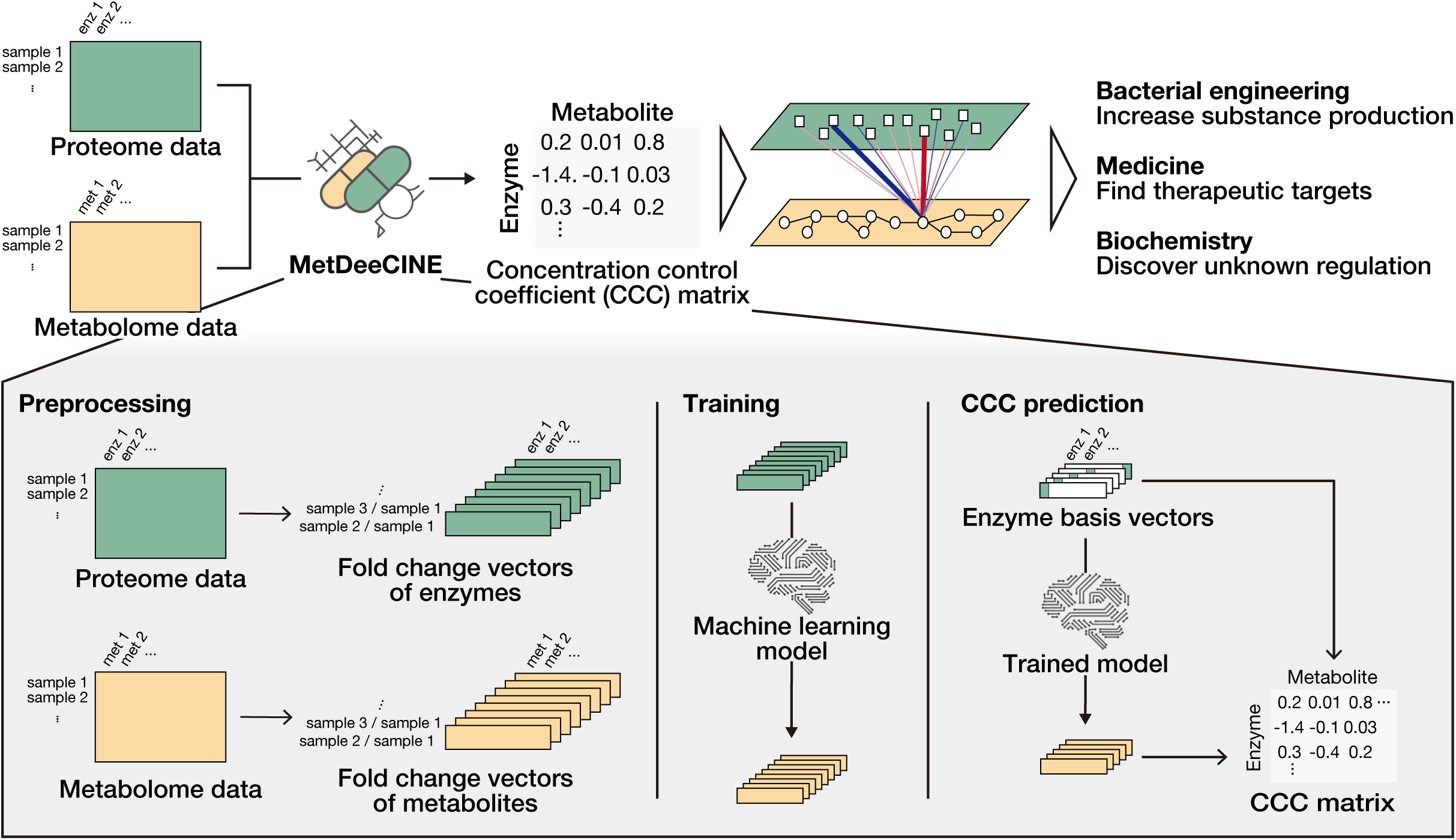
MetDeeCINE: A machine learning framework for predicting quantitative enzyme–metabolite relationships. MetDeeCINE is a machine learning framework that predicts the concentration control coefficient (CCC) matrix, as defined in metabolic control analysis (MCA), from multi-omics data (proteome and metabolome) (top). For each pair of samples, MetDeeCINE computes fold changes (FCs) in enzyme and metabolite concentrations (bottom left), trains an ML model that predicts metabolite FCs from enzyme FCs (bottom center), and then uses the trained model to simulate a scenario where only one enzyme changes, thereby constructing a CCC matrix (bottom right). The CCC matrix quantifies the effect of a change in each enzyme’s activity on the steady-state concentration of each metabolite.

## Results

### Metabolism-informed neural networks

Recent advances in the field of biology have led to the adoption of deep learning approaches that are guided by prior biological knowledge to handle sparsity in data and achieve better predictive performance, known as biology-informed neural networks (BINNs)^23–25^. Inspired by this approach, we hypothesized that incorporating information on metabolic reactions into a deep learning model would enhance its predictive accuracy. Although data such as detailed kinetic parameters and allosteric regulatory interactions are important, they are available only for certain model organisms (e.g., *E. coli* and yeast) or well-characterized metabolic pathways^26^. Heavy reliance on this detailed information makes it challenging to generalize the application of MetDeeCINE to less-annotated organisms or pathways. Moreover, providing excessive specific information could hinder the model’s ability to identify novel regulatory mechanisms hidden within the data.

To promote universal applicability, we propose “Metabolism-informed Neural Network (MiNN)” approaches that utilize only stoichiometric information, metabolites that serve as substrates, products, or cofactors for each enzymatic reaction. Such stoichiometric information is relatively easy to obtain or annotate for many species, making the model widely applicable and less constrained than the fully detailed kinetic approach. By mimicking metabolic processes and regulating model parameters based on reaction information, MiNN offers high interpretability and effectively incorporates prior biological knowledge.

In this study, we developed and evaluated two MiNN models. The first is the Metabolism-informed Bipartite Neural Network (MiBiNN) model, which treats the metabolic network as a bipartite graph composed of two classes of nodes — enzymes (reactions) and metabolites — connected in an alternating manner. MiBiNN directly models this flow of information by stacking layers of enzyme nodes and metabolite nodes in an alternating fashion (Fig. 2a, Methods). While MiBiNN explicitly represents the metabolic network structure in detail, our second approach, the Metabolism-informed Graph Neural Network (MiGNN), focuses on inter-metabolite relationships, resulting in a simpler model with fewer parameters and higher computational efficiency (Fig. 2b). MiGNN represents metabolites as nodes and connects them based on shared enzymatic reactions, allowing information to flow between metabolites that are directly or indirectly linked through the metabolic network.

**Figure 2.**
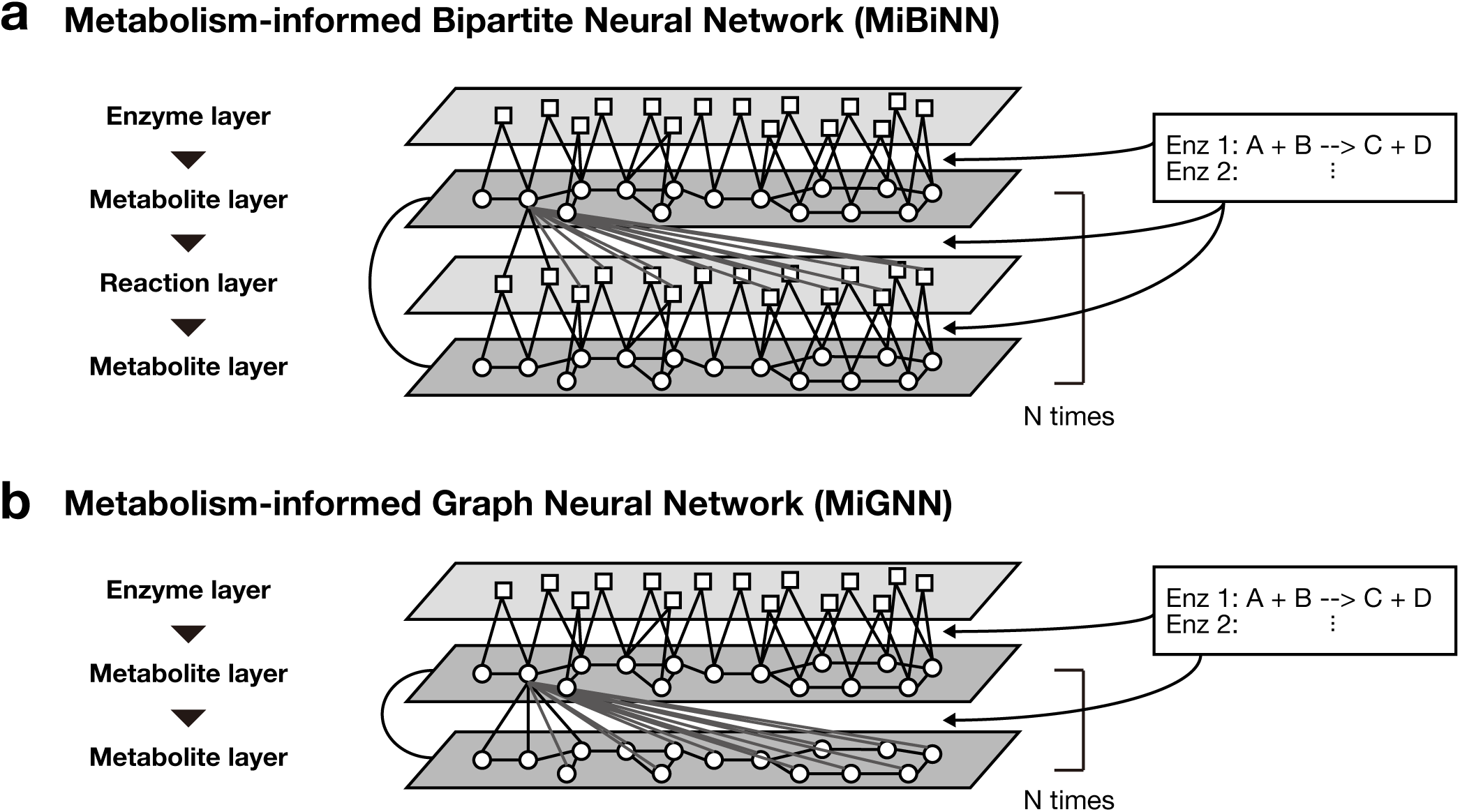
Metabolism-informed Neural Networks leverage stoichiometric information to improve interpretability and prediction accuracy. (a) Architecture of the Metabolism-informed Bipartite Neural Network (MiBiNN). MiBiNN is designed as a bipartite graph comprising alternating layers of reaction (enzyme) and metabolite nodes. Each reaction node is connected to the substrate, product, or cofactor metabolite nodes associated with that reaction, whereas each metabolite node is connected to all reaction nodes. To enforce this structure, large weights outside the substrate/product relationships are penalized in the loss function (see Methods). Residual connections link successive metabolite layers. This architecture mimics the flow of information in metabolic pathways. (b) Architecture of the Metabolism-informed Graph Neural Network (MiGNN). MiGNN omits the reaction layers directly representing metabolites as nodes and using edges to represent reactions between them. Metabolites appearing together in the same reaction are connected by an edge, whereas non-adjacent metabolites are also connected but connections between non-adjacent metabolites are penalized if their connections receive large weights. This design simplifies the network while retaining key information about metabolic connectivity.

### MiGNN outperforms other machine learning models in predicting CCCs derived from an *E. coli* kinetic model

To validate our approach, we evaluated the performance of MetDeeCINE on data and CCCs derived from an existing, small, yet well-curated kinetic model of *E. coli* central carbon metabolism^27^. Specifically, we used the default enzyme values in the *E. coli* kinetic model to generate simulated proteome data (by adding random perturbations to enzyme concentrations) and then performed steady-state simulations to obtain the corresponding metabolome profiles for MetDeeCINE training and evaluation (Fig. 3a).

**Figure 3.**
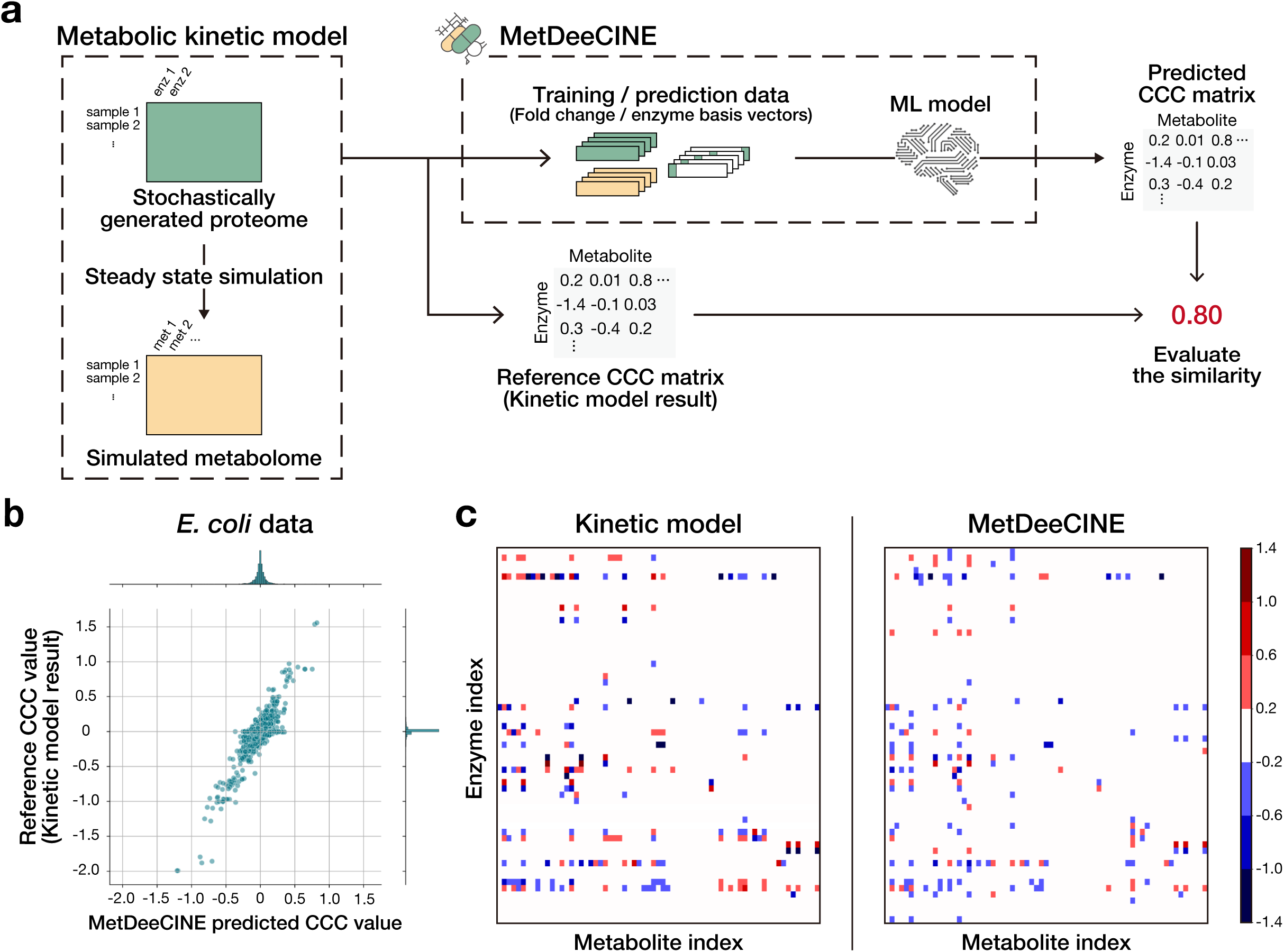
MetDeeCINE accurately predicts CCCs from an *E. coli* kinetic model. (a) Overview of the *in silico* experimental design using an existing *E. coli* kinetic model^27^. Enzyme concentrations were randomly perturbed to generate *in silico* proteome data. Steady-state simulations of the kinetic model were performed to generate the corresponding simulated metabolome data. These data were used to construct the training data and the reference CCC matrix for evaluation. We compared multiple machine learning models (Table 1) and selected MiGNN as the core component of MetDeeCINE. (b) Comparison of the reference and the predicted CCC matrices from MetDeeCINE (using MiGNN) for the *E. coli* kinetic model. Each point represents an element of the CCC matrix, quantifying the effect of small changes in enzyme activity on the steady-state metabolite concentrations for a specific enzyme-metabolite pair. The high correlation between predicted and reference CCCs demonstrates MetDeeCINE’s accuracy. (c) Heatmaps of the reference CCC matrix (left) and the CCC matrix predicted by MetDeeCINE (right). Columns represent the 67 metabolites and rows represent the 60 enzymes in the kinetic model. Colors indicate the CCC value for each enzyme-metabolite pair. The close similarity between the two heatmaps indicates that MetDeeCINE captures the overall regulatory structure of the metabolic network.

We trained and compared our new MiNN models, MiBiNN and MiGNN, alongside the commonly used machine learning models (linear regression and MLP) (Fig. S1, Methods). For each model, we optimized its architecture and hyperparameters (Table S1) based on the correlation between predicted and simulated metabolite FCs. Then, we used the best-trained model to predict the CCC matrix and compared the results with the reference CCCs from the kinetic model simulations. Among all the tested ML models, MiGNN achieved the best Spearman’s correlation (SCC) and lowest mean squared error (MSE) (Table 1). Although linear regression yielded the highest Pearson’s correlation (PCC), CCC values of the reference CCC matrix from the kinetic model exhibited a non-normal distribution (Fig. 3b), making SCC and MSE more appropriate metrics. Hence, MiGNN was chosen as the core component of MetDeeCINE for the following experiments. MiGNN’s predicted CCC matrix values closely matched the simulation-based ground truth (reference), capturing the overall regulatory structure of the model (Fig. 3b,c). These results indicate that MetDeeCINE can accurately infer metabolic regulation, even in the absence of detailed kinetic parameters or reaction-level information, highlighting its potential for broad applicability.

**Table 1.**
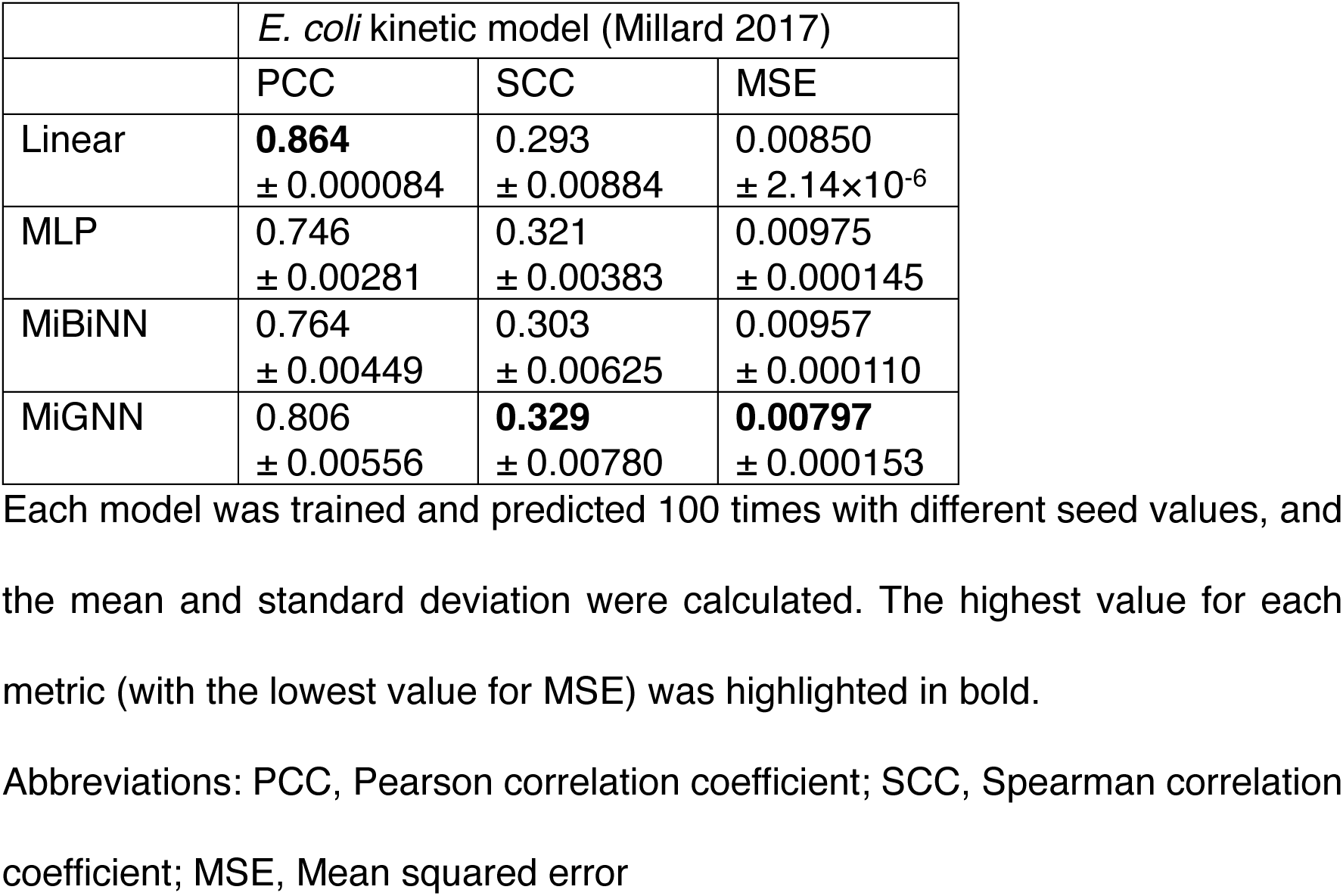
Performance metrics of each machine learning model.

|  | *E. coli* kinetic model (Millard 2017) |  |  |
| --- | --- | --- | --- |
|  | PCC | SCC | MSE |
| Linear | **0.864**<br>± 0.000084 | 0.293<br>± 0.00884 | 0.00850<br>± 2.14×10<sup>-6</sup> |
| MLP | 0.746<br>± 0.00281 | 0.321<br>± 0.00383 | 0.00975<br>± 0.000145 |
| MiBiNN | 0.764<br>± 0.00449 | 0.303<br>± 0.00625 | 0.00957<br>± 0.000110 |
| MiGNN | 0.806<br>± 0.00556 | **0.329**<br>± 0.00780 | **0.00797**<br>± 0.000153 |

Each model was trained and predicted 100 times with different seed values, and the mean and standard deviation were calculated. The highest value for each metric (with the lowest value for MSE) was highlighted in bold.
Abbreviations: PCC, Pearson correlation coefficient; SCC, Spearman correlation coefficient; MSE, Mean squared error

### MetDeeCINE accurately predicts complex enzyme–metabolite relationships in *E. coli*

Having confirmed that MiGNN-based MetDeeCINE can predict the entire CCC matrix with high accuracy, we next performed a more detailed evaluation of specific metabolites and enzymes to evaluate MetDeeCINE capabilities. We focused on two questions: (1) how accurately does MetDeeCINE predict which enzymes control the concentration of a specific metabolite and (2) how accurately does it predict how the change in the activity of a specific enzyme affects multiple metabolites across the network?

Pyruvate (PYR) is the end product of glycolysis that feeds into the TCA cycle via conversion to acetyl-CoA. It is also the starting point of various pathways including fermentation and amino acid biosynthesis, and is known to be abnormally regulated in cancer^28^. In the kinetic model, the time variation of pyruvate concentration is defined as being affected by multiple enzymes^27^, but its steady-state concentration is primarily affected only by pyruvate dehydrogenase (pdh) (Fig. 4a). MetDeeCINE predicted a value with large magnitude only for pdh CCC for pyruvate and all other enzymes’ CCCs were near zero (Fig. 4b), indicating that MetDeeCINE correctly identifies the primary enzyme controlling pyruvate concentration among various pyruvate-producing and -consuming enzymes.

**Figure 4.**
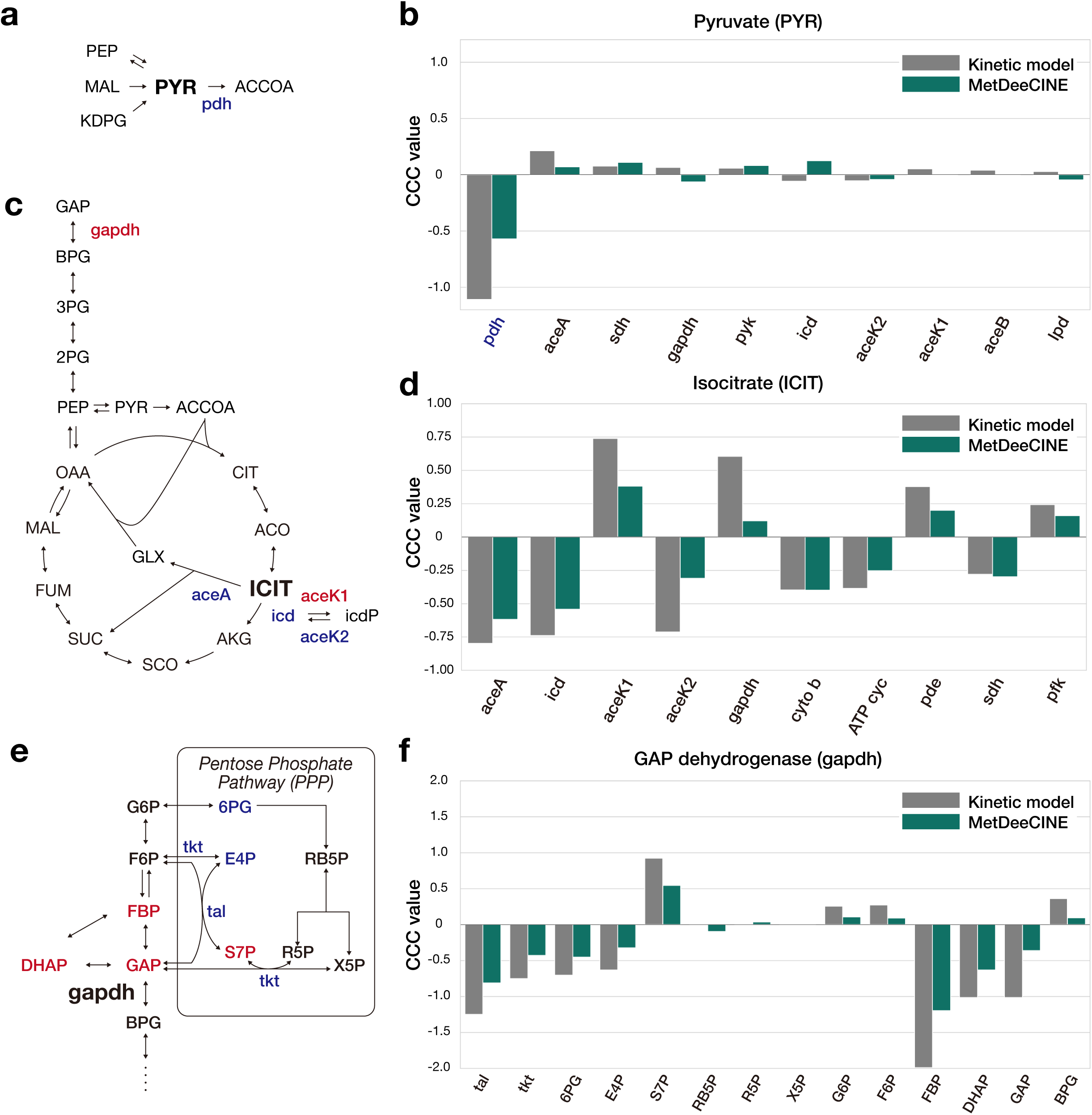
MetDeeCINE captures complex metabolic regulation, including allosteric effects, in *E. coli*. (a) Pyruvate (PYR)-related reactions in the *E. coli* kinetic model. Metabolites are shown in uppercase, and enzymes in lowercase. (b) Comparison of kinetic model-simulated (gray) and MetDeeCINE-predicted (green) CCC values for the top 10 enzymes with the greatest influence on pyruvate concentration. (c) Schematic representation of glycolysis and the TCA cycle in *E. coli*. Enzymes with large positive CCC values for isocitrate (ICIT) are shown in red, and those with negative CCC values with large magnitude are shown in blue. (d) Comparison of kinetic model-simulated (gray) and MetDeeCINE-predicted (green) CCC values for the top 10 enzymes with the greatest influence on isocitrate concentration. (e) Reactions involving glyceraldehyde-3-phosphate dehydrogenase (gapdh). Some PPP metabolites are influenced by gapdh activity, either positively (red) or negatively (blue). tal and tkt are enzymes, but their concentrations are altered by the addition of carbon, so they were also treated as metabolites in the analysis. (f) Comparison of kinetic model-simulated (gray) and MetDeeCINE-predicted (green) CCC values for gapdh with respect to metabolites highlighted in (e). MetDeeCINE yields values remarkably close to those from the kinetic model, despite using minimal prior knowledge. Abbreviations: 2PG, 2-phosphoglycerate; 3PG, 3-phosphoglycerate; 6PG, 6-phosphogluconate; aceK1, isocitrate dehydrogenase kinase; aceK2, isocitrate dehydrogenase phosphatase; aceA, isocitrate lyase; aceB, malate synthase; ACO, aconitate; ACCOA, acetyl-CoA; AKG, alpha-ketoglutarate; ATP cyc, ATP synthase; BPG, 1,3-bisphosphoglycerate; CIT, citrate; cyto b, cytochrome b; dbf, 2-dehydro-3-deoxy-phosphogluconate aldolase; DHAP, dihydroxyacetone phosphate; E4P, erythrose 4-phosphate; F6P, fructose-6-phosphate; FBP, fructose 1,6-bisphosphate; FUM, fumarate; G6P, glucose-6-phosphate; GAP, glyceraldehyde-3-phosphate; gapdh, glyceraldehyde-3-phosphate dehydrogenase; GLX, glyoxylate; icd, isocitrate dehydrogenase; icdP, phosphorylated isocitrate dehydrogenase; ICIT, isocitrate; KDPG, 2-keto-3-deoxy-6-phosphogluconate; lpd, dihydrolipoamide dehydrogenase; MAL, malate; OAA, oxaloacetate; pdh, pyruvate dehydrogenase complex; pde, phosphodiesterase; pdk, pyruvate dehydrogenase kinase; pfk, phosphofructokinase; PEP, phosphoenolpyruvate; pyk, pyruvate kinase; PYR, pyruvate; R5P, ribose 5-phosphate; RB5P, ribulose 5-phosphate; SCO, succinyl-CoA; SDH, succinate dehydrogenase; S7P, sedoheptulose 7-phosphate; SUC, succinate; tal, transaldolase; tkt, transketolase; X5P, xylulose 5-phosphate

Another TCA cycle metabolite, isocitrate (ICIT), is catalytically produced by aconitase 2 and consumed by isocitrate dehydrogenase, but owing to the effects of cofactors and allosteric control, its steady-state concentration in the kinetic model is further regulated by kinase/phosphatase for isocitrate dehydrogenase, the glycolytic enzyme glyceraldehyde-3-phosphate dehydrogenase (gapdh), and enzymes involved in the electron transport chain or ATP synthesis/consumption^27^ (Fig. 4c). MetDeeCINE accurately predicted not only the direction (positive or negative) but also the magnitude of each enzyme’s influence on isocitrate (Fig. 4d), suggesting that MetDeeCINE captures allosteric effects that are not explicitly provided in the stoichiometric information alone.

We then examined whether MetDeeCINE could accurately predict how a change in the activity of a single enzyme influences multiple metabolites. The glycolytic enzyme gapdh is known to affect metabolites in the pentose phosphate pathway (PPP)^27^. In the kinetic model, the CCC of gapdh with respect to PPP metabolites showed a highly complex pattern: strongly negative values for some metabolites, a distinctly large positive value only for sedoheptulose 7-phosphate (S7P), and near-zero values for others (Fig. 4e). MetDeeCINE again reproduced this complex pattern with high fidelity (Fig. 4f).

Collectively, these findings demonstrate that MetDeeCINE accurately predicts enzyme–metabolite relationships across the entire metabolic network. Notably, the ability to capture the impact of allosteric regulation (e.g., the effect of gapdh on isocitrate in the TCA cycle) suggests that MetDeeCINE not only incorporates stoichiometric information but also learns regulatory mechanisms embedded within the data. Moreover, by accurately predicting how the perturbation of a single enzyme affects multiple metabolites, MetDeeCINE can yield insights into detailed regulatory relationships that are often overlooked in single-omics analyses.

### MetDeeCINE identifies the rate-limiting enzyme of gluconeogenesis in mouse *in vivo* data

While MiGNN performed well with *E. coli* data, mammalian metabolism is more complex, and *in vivo* experimental data can exhibit more intricate behavior than well-defined kinetic-model simulations. To test the utility of MetDeeCINE in a mammalian *in vivo* setting, we applied it to a multi-omics dataset of mouse livers that we previously reported^14^. This dataset contains transcriptome, proteome, and metabolome measurements (n=47) from wild-type (WT) and leptin-deficient *ob/ob* mice under two conditions: after 16 hours of fasting and 4 hours after an oral glucose challenge (Fig. 5a). As a proof-of-concept, and to be consistent with our previous work^14^, we focused on enzymes and metabolites involved in central carbon metabolism, particularly the gluconeogenesis pathway. After hyperparameter tuning, the model’s final predictions for metabolite FCs showed a strong correlation with the experimental measurements (Fig. 5b).

**Figure 5.**
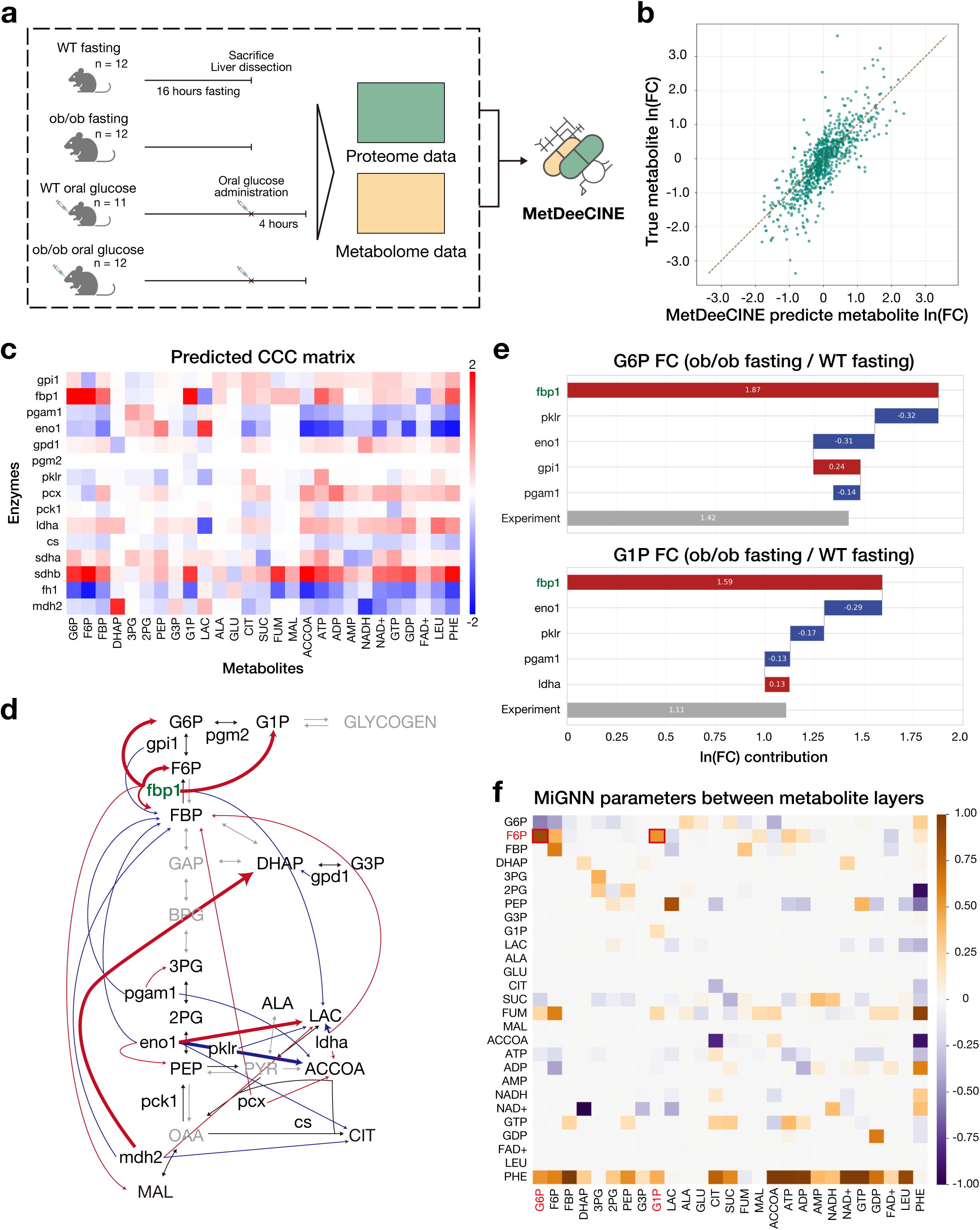
MetDeeCINE identified the rate-limiting enzyme of glycogenesis from mouse multi-omics data. (a) Workflow of the MetDeeCINE analysis using the mouse liver dataset. Wild-type and *ob/ob* (genetically obese) mice were sacrificed after 16 hours of fasting or 4 hours after an oral glucose challenge. Proteome and metabolome data from the liver were used to predict the CCC matrix. (b) Scatter plot of predicted (x-axis) versus observed (y-axis) metabolite FCs. Each point represents one element in the logarithm (log) FC vector for a specific metabolite. (c) Heatmap of the predicted CCC matrix for enzymes and metabolites involved in mouse gluconeogenesis. (d) Predicted enzyme–metabolite relationships (absolute value of CCC ≥ 0.5) mapped onto the gluconeogenesis pathway. Arrow thickness reflects the magnitude of the CCC value (0.5–1.0, 1.0–2.0, ≥2.0). Gray metabolites and enzymes (arrows) were not measured. fbp1 (green) is a known rate-limiting enzyme in gluconeogenesis. (e) Waterfall plot comparing MetDeeCINE-predicted contributions of individual enzymes with the observed log FC of glucose-6-phosphate (G6P, top) or glucose-1-phosphate (G1P, bottom) between fasted *ob/ob* mice and fasted WT mice (“Experiment”). (f) Heatmap of the metabolite-layer weight matrix from MiGNN. The connections (edges) between fructose-6-phosphate (F6P) and G6P or G1P (red box) are relatively strong, reflecting the known pathway structure in which fbp1 controls F6P, which in turn influences G6P and G1P. Abbreviations: 2PG, 2-phosphoglycerate; 3PG, 3-phosphoglycerate; ACCOA, acetyl-coa; ADP, adenosine diphosphate; ALA, alanine; AMP, adenosine monophosphate; ATP, adenosine triphosphate; CIT, citrate; cs, citrate synthase; DHAP, dihydroxyacetone phosphate; eno1, enolase 1; F1,6P, fructose 1,6-bisphosphate; F6P, fructose 6-phosphate; fbp1, fructose-1,6-bisphosphatase 1; FAD+, flavin adenine dinucleotide; fh1, fumarase 1; G1P, glucose 1-phosphate; G3P, glycerol 3-phosphate; G6P, glucose 6-phosphate; GDP, guanosine diphosphate; GLU, glutamate; gpd1, glycerol-3-phosphate dehydrogenase 1; gpi1, glucose phosphate isomerase 1; GTP, guanosine triphosphate; LAC, lactate; ldha, lactate dehydrogenase a; LEU, leucine; MAL, malate; mdh2, malate dehydrogenase 2; NAD+, nicotinamide adenine dinucleotide; NADH, nicotinamide adenine dinucleotide, reduced; pcx, pyruvate carboxylase; pck1, phosphoenolpyruvate carboxykinase 1; PEP, phosphoenolpyruvate; PHE, phenylalanine; pgam1, phosphoglycerate mutase 1; pgm2, phosphoglucomutase 2; pklr, pyruvate kinase, liver and red blood cell; sdha, succinate dehydrogenase complex subunit a; sdhb, succinate dehydrogenase complex subunit b; SUC, succinate

We then predicted the CCC matrix and mapped enzyme–metabolite pairs with absolute CCC values above 0.5 onto the metabolic pathway diagram (Fig. 5c,d). A major strength of MetDeeCINE is that it can be used for data with unmeasured enzymes or metabolites, which is difficult in many traditional approaches. The predicted CCC matrix highlights fructose-1,6-bisphosphatase 1 (fbp1) as having a strong influence on many metabolites (Fig. 5d), which is consistent with the known role of fbp1 as a rate-limiting enzyme in gluconeogenesis^29^.

In analyses of disease-related metabolic alterations, identifying which enzyme changes drive metabolite changes is often difficult because multiple enzymes can shift simultaneously, influencing one another and finally resulting in altered metabolite concentrations^14^. In our previous comparison of *ob/ob* mice (fasted) and WT mice (fasted), we observed an increase in glucose, glucose-6-phosphate (G6P), and glucose-1-phosphate (G1P), which are the end products of gluconeogenesis; however, the proteome data showed an increased abundance of multiple enzymes in the gluconeogenesis pathway (gpi1, pgam1, eno1, etc.) (Fig. S2). Hence, it was challenging to assess the individual contribution of each enzyme using a single-omics approach; therefore, we used MetDeeCINE to determine which enzyme had the greatest impact on the accumulation of G6P and G1P. MetDeeCINE predicted that fbp1, the rate-limiting gluconeogenic enzyme, is the primary contributor to increased G1P concentration. This prediction aligns with previously reported experimental evidence^29^ and further provides a quantitative explanation of how other enzymes influence G6P and G1P levels (Fig. 5e).

Moreover, analysis of the learned inter-metabolite weights in MiGNN revealed large connection weights between fructose-6-phosphate (F6P), the product of fbp1, and G6P or G1P (Fig. 5f). This implies that MetDeeCINE is not merely learning correlations, but also producing predictions consistent with known pathway structures, in which fbp1 alters F6P concentration, which in turn affects G6P and G1P. Hence, even with complex *in vivo* data from mice, MetDeeCINE generates biologically consistent results, demonstrating its ability to identify rate-limiting enzymes and contributors to abnormal metabolite accumulation, along with potential underlying mechanism, without requiring the detailed reaction reversibility or other constraints typically needed for kinetic models.

## Discussion

In this study, we developed MetDeeCINE, a novel machine learning framework for quantitatively and comprehensively analyzing metabolic regulation networks from multi-omics data. Previous approaches for multi-omics integration in metabolism have often required extensive information on each reaction, such as metabolic flux, kinetic parameters, or allosteric regulation, thereby limiting their applicability to well-annotated pathways or species. Consequently, many previous metabolic multi-omics studies have focused on analyzing individual omics layers separately. In contrast, MetDeeCINE predicts the CCC matrix — representing how changes in enzyme activity influence metabolite concentrations — using only the measured data and readily available stoichiometric information of metabolic reactions. This expands the applicability even to pathways or organisms for which detailed annotations are incomplete. We demonstrated that, in simulations with a kinetic model, our newly developed MiGNN had the best predictive performance among several machine learning methods, accurately reproducing the influence of an enzyme on not only its immediate substrates and products, but also on distant metabolites. Furthermore, applying MetDeeCINE to experimentally obtained mouse data produced biologically consistent predictions, revealing a rate-limiting enzyme in gluconeogenesis and identifying the enzymes responsible for G6P and G1P accumulation.

MiGNN, the core of MetDeeCINE, is a graph neural network designed to mirror the structure of metabolic networks, offering interpretability that helps overcome the “black box” issue often encountered in ML-based metabolic analyses. Beyond simply identifying important features, MiGNN’s weight matrices can be examined to elucidate the mechanistic basis underlying the effects of those features. For example, after confirming through CCC analysis that fbp1 is the primary rate-limiting enzyme in gluconeogenesis, we examined the metabolite-layer weight matrix and observed strong connections between F6P (produced by fbp1) and G6P or G1P, demonstrating that MiGNN’s prediction aligns with the known metabolic pathway. Moreover, our *E. coli* kinetic model experiment showed that MiGNN could capture the impact of allosteric regulation not encoded in the stoichiometric data alone (e.g., the effect of gapdh on isocitrate). The presence of a strong edge between two metabolites that are distant in the network suggests direct or cofactor-mediated allosteric regulation. Future work should include a detailed analysis of MiGNN weight matrices in the presence of known allosteric controls, potentially revealing unknown regulatory mechanisms that may indicate novel drug targets in less-studied metabolic pathways.

These results highlight a key advantage of MetDeeCINE over traditional approaches. While methods like flux balance analysis (FBA) are useful for predicting metabolic fluxes under steady-state conditions, they do not directly address the dynamic control of metabolite concentrations. Kinetic models, while capable of capturing dynamic behavior, are limited by their reliance on detailed ODEs and various parameters, which are often unavailable. MetDeeCINE, by contrast, leverages the strengths of both approaches: it incorporates the structural information of metabolic networks (like FBA) and utilizes data-driven learning to infer quantitative relationships between enzymes and metabolites (similar to kinetic models), but without requiring complete kinetic information.

MiGNN imposes constraints on the model structure based on metabolic mechanisms (e.g., partial connectivity between enzymes and metabolites, residual connections between metabolite layers, and stoichiometry-based regularization), helping the model avoid local minima and converge on biologically meaningful solutions. Notably, within the MetDeeCINE framework, the ML model is not directly trained on CCC values. Indeed, the CCC is computed from inputs that are distant, in terms of Euclidean distance, from those used in training. This approach makes the predicted CCC matrix susceptible to overfitting, thereby rendering metabolism-specific constraints particularly effective. On the other hand, MiBiNN — another MiNN approach — did not outperform MiGNN or standard ML models. This is likely due to its larger number of parameters, which complicated the training process, and/or overly strong constraints that limited model flexibility.

Compared to the reference CCC matrix from the kinetic model, predicted CCC matrix for the experimental dataset were denser and exhibited larger values. In kinetic models, the defined system is “closed” within the model’s enzymes and metabolites, whereas real biological data are subject to many unmeasured influences. Since our ML models attempt to explain metabolite variation using only the measured enzyme FCs, it may overfit and generate inflated CCC values. Another potential source of overfitting stems from the need to predict metabolite changes that are nearly zero between experimental conditions. Addressing missing enzymes and metabolites in the pathway, as well as focusing on highly variable enzymes and metabolites, may improve MetDeeCINE’s performance in the future.

There are several limitations to our framework. First, the current model cannot account for the impact of unmeasured enzymes on metabolite fluctuations, nor can it estimate the uncertainty of predictions stemming from these unmeasured variables. Future extensions could incorporate probabilistic modeling or ensemble methods to address this limitation. Second, the present version treats regulation as unidirectional (enzyme → metabolite), despite the fact that actual metabolism involves feedback from metabolites to enzymes (e.g., allosteric regulation). Although our results suggest that MiGNN can implicitly capture some aspects of allosteric regulation, future versions could explicitly incorporate feedback loops. Third, while CCCs formally represent the impact of enzyme activity, we used measured enzyme abundance as a proxy for activity, thus neglecting the kinetic parameters that also influence activity. In addition, MetDeeCINE’s predictions rely on a local linear approximation (small fold change), although we observed better predictive performance with larger FCs. Potential solutions include partially excluding certain enzymes from the proteome to quantify uncertainty, substituting proteome data with transcriptome data, and developing new methods for a more detailed analysis of MiGNN edges and predicted CCCs.

Despite these challenges, MetDeeCINE’s integration of measured data and readily available stoichiometric information offers a scalable and biologically valid deep learning approach applicable even when annotations are sparse. It substantially enhances the resolution of metabolic regulation studies compared to single-omics approaches. Applying MetDeeCINE to disease datasets can identify enzymes driving abnormal metabolite accumulation, thus highlighting potential therapeutic targets. In conclusion, MetDeeCINE represents a significant step towards a comprehensive, systems-level understanding of metabolic regulation, bridging the gap between detailed kinetic modeling and data-driven machine learning approaches. We anticipate that MetDeeCINE will be a valuable tool for researchers in a wide range of fields, from basic biology to medicine and biotechnology.

## Materials and Methods

### Study overview

In this study, we developed MetDeeCINE, a novel framework that predicts concentration control coefficients (CCCs) from multi-omics data, focusing on the quantitative relationship between enzymes and metabolites. CCCs, as defined by Metabolic Control Analysis (MCA)^8^, measure how small changes in enzyme activities in a steady-state condition affect metabolite concentrations. Crucially, CCCs capture both direct (i.e., the metabolite that is directly produced or consumed by the enzymatic reaction) and indirect relationships mediated through multiple reactions and metabolites. While MCA often focuses on flux control coefficients, investigating the control of steady-state metabolite concentrations is also crucial, particularly in light of oncometabolite biology in cancer and the optimization of compound production in metabolic engineering. MetDeeCINE uses proteome and metabolome data obtained from multiple samples (organisms, conditions) to compute fold changes (FCs) between sample pairs. It then trains a machine learning (ML) model that takes enzyme FC vectors as input and predicts metabolite FC vectors. Once trained, the model can be used to estimate CCCs without requiring detailed kinetic parameters (Km, kcat, etc.) or differential equations. Consequently, MetDeeCINE provides a data-driven method for deciphering quantitative relationships between enzymes and metabolites at an omics-scale.

### Kinetic model of metabolism

We used the *E. coli* central carbon metabolism model developed by Millard *et al.*^27^ for the kinetic model of metabolism. This model comprises 62 metabolites and 68 metabolic and transport reactions. Many enzymes within the model are treated as having fixed concentrations, while some enzyme concentrations are altered by phosphorylation or the addition of carbon groups. In our definition, components with changing concentrations were classified as metabolites, whereas other reaction-mediating components were classified as enzymes. According to this definition, the model contained 60 enzymes and 67 metabolites. Default values were used for all initial metabolite concentrations and enzyme kinetic parameters. Simulations were performed in MATLAB (version 9.14) using SimBiology (version 6.4.1).

### Generation of in silico multi-omics data from the metabolic kinetic model

We performed multiple steady-state simulations using the kinetic model, with random perturbations applied to enzyme concentrations. This generated a set of simulated proteome data (enzyme concentrations) and corresponding metabolomic data (metabolite concentrations). We introduced two types of perturbations: first, we created three “strain enzyme vectors” by adding a large perturbation to the default enzyme concentrations in the kinetic model (WT enzyme vector). These large perturbations correspond to differences in strains and experimental conditions. The enzyme concentration for strain X, 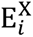, obtained by perturbing the default enzyme concentration E_*i*_ for enzyme *i*, is defined as follows:

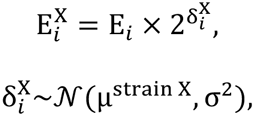

where µ^strain 1^ = 0.5, µ^strain 2^ = −0.5, µ^strain 3^ = 1 and σ^2^ = 0.3^2^.

We then generated “sample enzyme vectors” by applying small perturbations to the WT and strain enzyme vectors. These small perturbations corresponded to individual differences; 50 sample enzyme vectors were generated from each strain vector. *i*-th element of the *a*-th sample enzyme vector of strain X was denoted 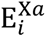 and determined according to the following equation:

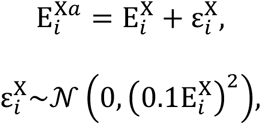

and sample enzyme vectors from WT enzyme vectors were generated in the same manner. By conducting a steady-state analysis of the kinetic model with each sample enzyme vector as the initial values for the enzyme concentrations, we generated corresponding sample metabolite vector, representing the metabolite concentrations at steady state. Any sample enzyme vector that failed to reach a steady state was discarded and regenerated. Finally, we added random noise to each sample enzyme and metabolite vector to simulate measurement error. The concentration of enzyme *i* in sample a, 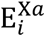, and the concentration of metabolite *j*, 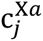, were adjusted by adding noise according to the following formula:

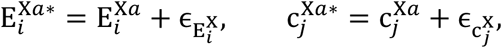

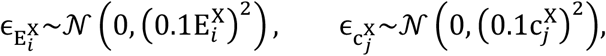

where 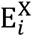 and 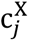 are the values of the strain vectors of strain X corresponding to enzyme *i* and metabolite *j*, respectively. Since the perturbations corresponding to differences in experimental conditions and individual differences are added to the enzyme vector, the resulting metabolite concentrations obtained by inputting these perturbed enzyme vectors into the kinetic model and enzyme concentrations satisfy the steady-state conditions. However, the addition of noise disrupts this steady state. As a result, we obtained 50 samples each for wild type (WT) and three different strains, yielding enzyme and metabolite concentration vectors for each sample.

### Computing the reference CCC matrix

In MCA, the concentration control coefficient is a quantitative measure of the impact of changes in enzyme activities on changes in metabolite levels, and it is defined as 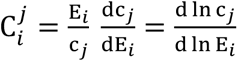, where ln denotes the natural logarithm. We calculated the reference CCC using the kinetic model via:

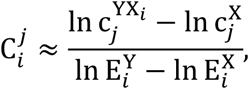

where 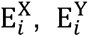, are the concentrations of enzyme *i* in strain X and strain Y, 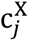 is the steady-state concentration of metabolite *j* in strain X, and 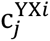 is the steady-state concentration of metabolite *j* obtained by substituting only enzyme *i* in strain X with its value in strain Y.

### Experimental data acquisition and preprocessing

We obtained proteome and metabolome data for the mouse liver from our previous study^14^. In brief, wild-type (WT) and leptin-deficient (*ob/ob*) C57BL/6J male mice (10 weeks old) were subjected to two conditions: the fasting group, sacrificed after 16 hours of fasting, and the oral glucose group, which received glucose orally (2 g glucose per kg body weight) after fasting and was sacrificed 4 hours later. Proteome and metabolome data were preprocessed as previously described^14^. Missing values in the enzyme measurements were imputed using the mean value for each experimental condition. Enzymes that were missing in all samples of a particular condition were excluded from the analysis. For metabolites, missing values were not imputed, and any missing measurements for a particular metabolite were just excluded from the loss function in MetDeeCINE.

### MetDeeCINE

MetDeeCINE consists of three steps: (1) preprocessing the enzyme and metabolite concentration vectors for each individual (sample), (2) training a machine learning model, and (3) predicting the CCC matrix using the trained model. As a preprocessing step, the natural logarithm (log) fold change **X** and **M**^true^ between two sample pairs were calculated separately for enzyme and metabolite concentration vectors, and used as training data for the machine learning model. We examined two methods for calculating FCs: using all pairs of samples, and using all pairs except for those with the same experimental conditions. The machine learning models predict metabolite FC vectors from enzyme FC vectors. We developed two novel metabolism-informed neural networks, MiBiNN and MiGNN, and evaluated their performance in predicting the CCC matrix.

### MiBiNN

The Metabolism-informed Bipartite Neural Network (MiBiNN) is designed based on the principle that a metabolic network can be represented as a bipartite graph consisting of enzymes (reactions) and metabolites. Each reaction converts specific metabolites (substrates and cofactors) into products, influencing the reaction rates of other reactions through changes in substrate or allosteric regulator concentrations. MiBiNN reflects this by alternating between reaction and metabolite layers while sharing the same parameters in repetition, as described in the following equation:

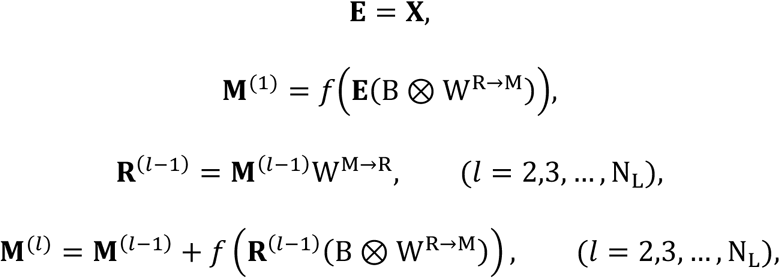

where **X** is the natural log fold change in enzyme abundance between the two samples, 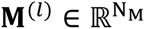 and 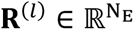 are the *l*-th metabolite and reaction layer, respectively, and *f* is the activation function. 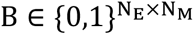 (N_E_ and N_M_ are the number of enzymes and metabolites, respectively) is a mask matrix for the learnable parameter W, where B_*ij*_ is 1 if metabolite *j* is a substrate or product of the reaction catalyzed by enzyme *i*, and 0 otherwise. B was defined from the stoichiometric matrix of the kinetic model when learning kinetic model-derived data, and from the KEGG^30^ database when learning experimental data by obtaining the substrate or product of the reaction associated with each enzyme. This structure ensures that each node in the enzyme and reaction layers is connected only to the metabolite nodes corresponding to the substrate, products, and cofactors of the associated reaction. Conversely, each node in the metabolite layer is connected to all reaction nodes. The loss function ℒ_MiBiNN_ for MiBiNN is expressed as follows:

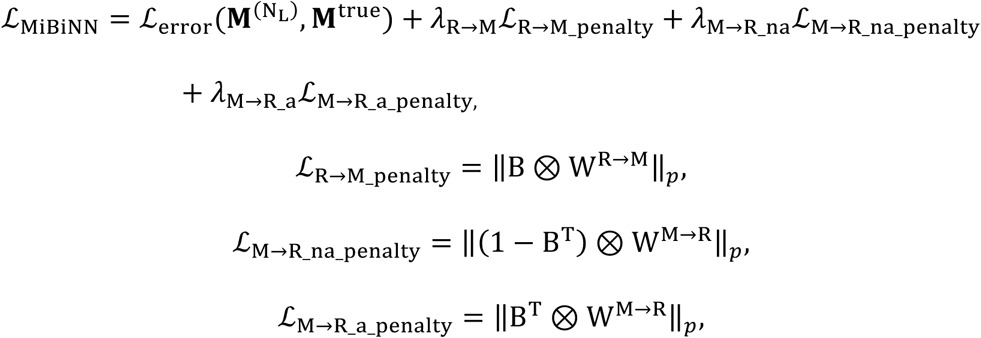

where 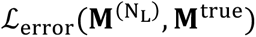 is the error function between the true data and predictions. ℒ_R→M_penalty_, ℒ_M→R_na_penalty_ and ℒ_M→R_a_penalty_ are the loss functions corresponding to edges from each enzyme (reaction) node to the substrate, product, cofactor node, edges of each metabolite node to reaction nodes that are not the formation/degradation reaction, edges of each metabolite node to the formation/degradation reaction node, and *λ*_R→M_, *λ*_M→R_na_ and *λ*_M→R_a_ are the corresponding regularization parameters. ‖C‖_*p*_ (*p* ∈ {1, 2}) stands for 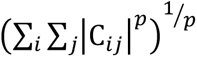. By setting *λ*_M→R_a_ smaller than *λ*_M→R_na_, regularization terms keep weights corresponding to non-biological edges small while allowing edges consistent with stoichiometric relationships to be learned more freely. And yet if the value of some edge that does not follow the stoichiometry is large, it suggests the presence of a control such as an allosteric control. Thus, MiBiNN incorporates the stoichiometry of metabolic reactions into both the connectivity between nodes in the neural network and the regularization terms in the loss function, resulting in a deep learning model that reflects the underlying mechanisms of metabolism.

### MiGNN

MiGNN is a simplified model derived by eliminating the reaction layers present between the metabolite layers. Instead, reactions connecting the metabolites are represented by inter-metabolite edges. MiGNN can be written as follows:

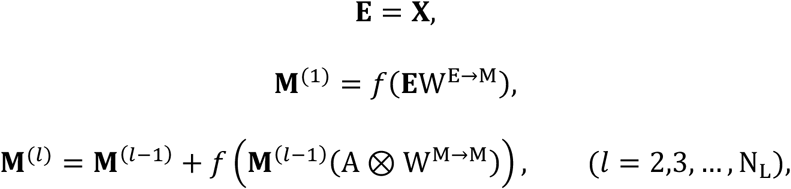

and the loss function ℒ_MiGNN_ is expressed as follows:

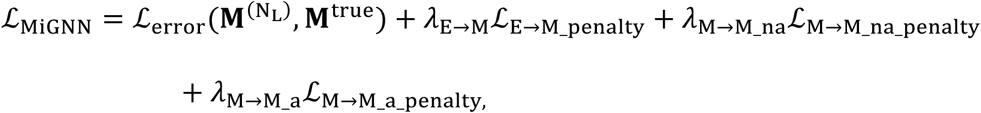

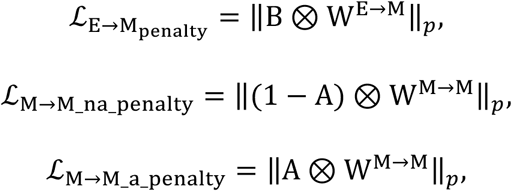

where 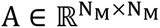 is the adjacency matrix, A_*ij*_ is 1 if metabolite *i* and metabolite *j* are produced or consumed in the same reaction, and 0 otherwise. ℒ_E→M_penalty_, ℒ_M→M_na_penalty_ and ℒ_M→M_a_penalty_ are regularization terms respectively for the edges from each enzyme (reaction) node to the substrate, product, and cofactor nodes, between each metabolite node with zero adjacency matrix values, and between each metabolite node with one adjacency matrix value, where *λ*_E→M_, *λ*_M→M_na_ and *λ*_M→M_a_ratio_ are the corresponding regularization parameters. In other words, each node in the enzyme layer is connected to the metabolite nodes representing the substrates, products, and cofactors of the corresponding reaction. Each node in the metabolite layers is connected to all other metabolite nodes. However, similar to MiBiNN, MiGNN applies regularization terms that discourage large weights on edges not supported by stoichiometry.

### Predicting the CCC matrix

The CCC matrix is constructed using the trained model by inputting a set of “enzyme basis vectors”. Each of these vectors has only one non-zero element (equal to the natural log FC of a specific enzyme between two conditions), while all other elements are zero. The *ij* component 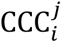 of the CCC matrix is calculated as follows:

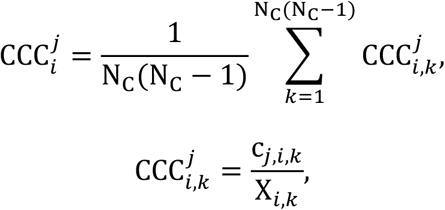

where X_*i*,*k*_ is the natural log FC of enzyme *i* for sample pair *k* (*k* = 1,2, …, N_L_(N_L_ − 1)) (N_L_ is the number of experimental conditions). c_j,*i*,*k*_ is the model output when inputting the *i*-th enzyme basis vector, in which the value of the *i*-th dimension is equal to the natural log FC of enzyme *i* and the values of other dimensions are 0 (= ln(1)). If the proteome data contain N_E_ enzymes, N_E_N_L_(N_L_ − 1) enzyme basis vectors are created. CCC in MCA can be expressed as 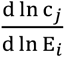, where E is the enzyme activity and c_j_ is the metabolite concentration, each element in the CCC matrix calculated by MetDeeCINE can be interpreted as a finite-difference approximation of this definition.

### Hyperparameter optimization

We used Optuna^31^ to optimize the hyperparameters listed in Table S1, with 500 trials for MiBiNN, MiGNN, and MLP, and 200 for linear regression. The evaluation metric was the mean Pearson correlation coefficient (PCC) between predicted and actual metabolite FCs on the validation data, using a cross-validation setup. Specifically, we used condition-based splitting: for fold *i*, FCs that contained condition *i* samples were used as validation data, while other data were used as training data. This approach prevented the training data from containing samples that were highly similar to the validation data, thus providing a robust evaluation of the model’s generalization performance. The early stopping patience was set to 10, the learning rate was 0.001, the batch size was 512 for kinetic model data and 32 for experimental data. We used AdamW for optimization. Model construction and training were implemented in PyTorch (version 2.0.1).

### Calculating individual enzyme contributions to metabolite changes

To quantify the contribution of each each enzyme to the change in a metabolite under specific conditions, we multiplied the corresponding MetDeeCINE-predicted CCC value by the average log FC of that enzyme across those conditions. We then compared the sum of these predicted contributions to the experimentally observed log FC of the metabolite.

## ACKNOWLEDGEMENTS

This work was supported by KAKENHI grants from the Japan Society for the Promotion of Science (JSPS) to S.O. and H.S (24K03030). We thank all the laboratory members for discussion and K. Tanaka for help with preparation of the manuscript.

## AUTHOR CONTRIBUTIONS

S.O. conceived of and designed the study. T.I. developed MetDeeCINE, performed computational study with inputs from S.O., Y.W. and S.U. T.I., S.O., and H.S. wrote the manuscript. S.K. and H.S. supervised the study.

## Supplementary Information

**Supplementary Table 1.** Hyperparameter options and selections.

|  |  |  |
| --- | --- | --- |
|  |  | E. coli kinetic model (Millard 2016) |
| **ML architecture** | Linear / MLP / MiBiNN / MiGNN | MiGNN |
| **Preprocessing** |  |  |
| FC calculation | Between all samples/<br>Exclude within the same group | Exclude within the same group |
| **Model property** |  |  |
| Number of layers (except linear model) | 1 – 4 | 2 |
| Fit loss function | MSE / MAE | MSE |
| Regularization type | L1 / L2 | L2 |
| Regularization strength |  | 1×10<sup>-5</sup> for enzyme-metabolite edges and associated metabolite-metabolite edges<br>1×10<sup>-3</sup> for unassociated metabolite-metabolite edges |
| Activation function (except linear model) | ReLU / tanh / elu / mish / swish | tanh |

**Supplementary Figure 1.**
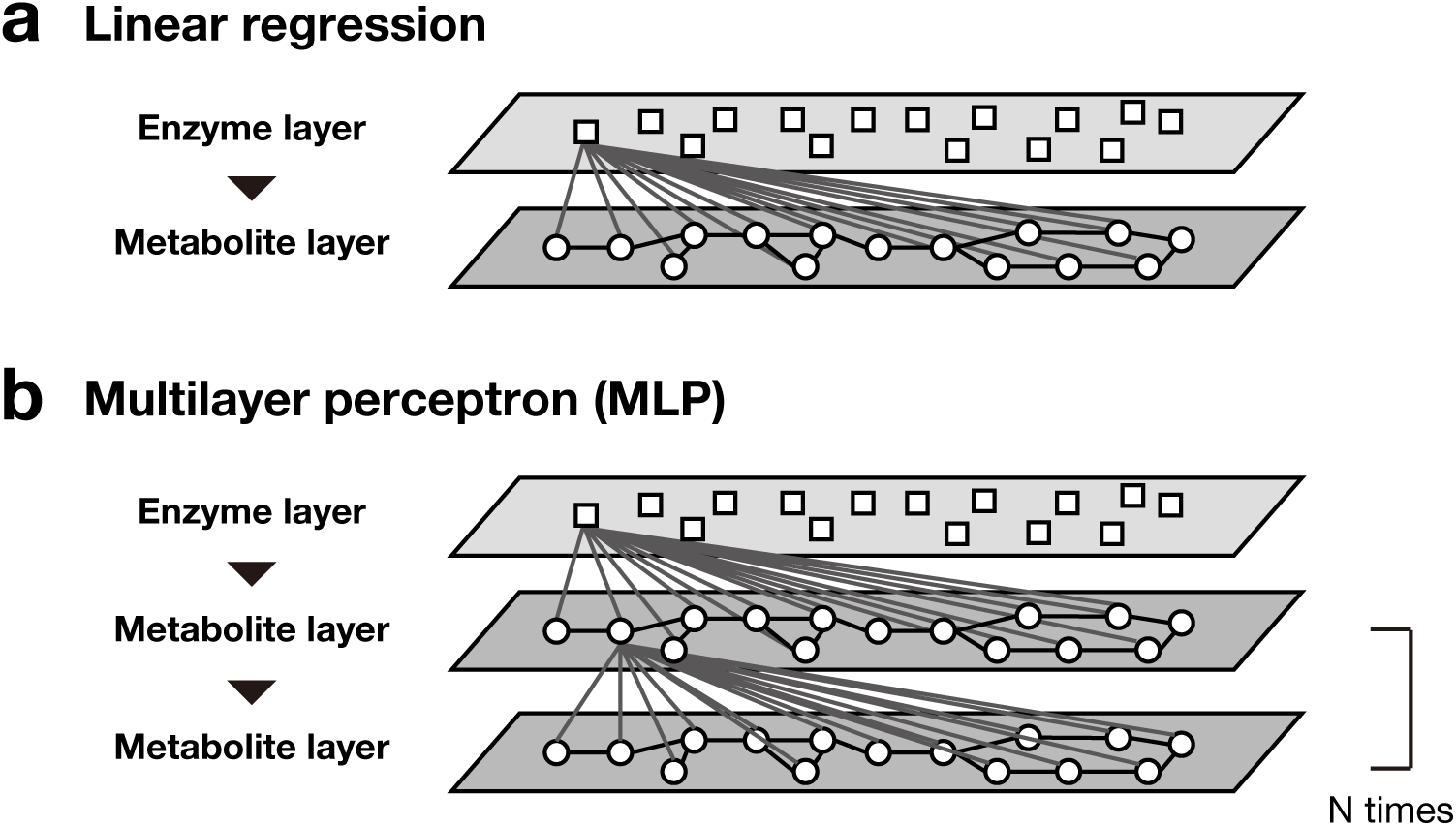
Architecture of classical machine learning models. (a) Structure of the linear regression model. It consists of an input layer for enzyme FCs and an output layer for metabolite FCs, where the value of each metabolite node is calculated as the linear combination of the values of all enzyme nodes. (b) Structure of multilayer perceptron (MLP). The number of nodes in the hidden layer was set equal to the number of metabolites. All layers are fully connected, with each node receiving input from all nodes in the preceding layer.

**Supplementary Figure 2.**
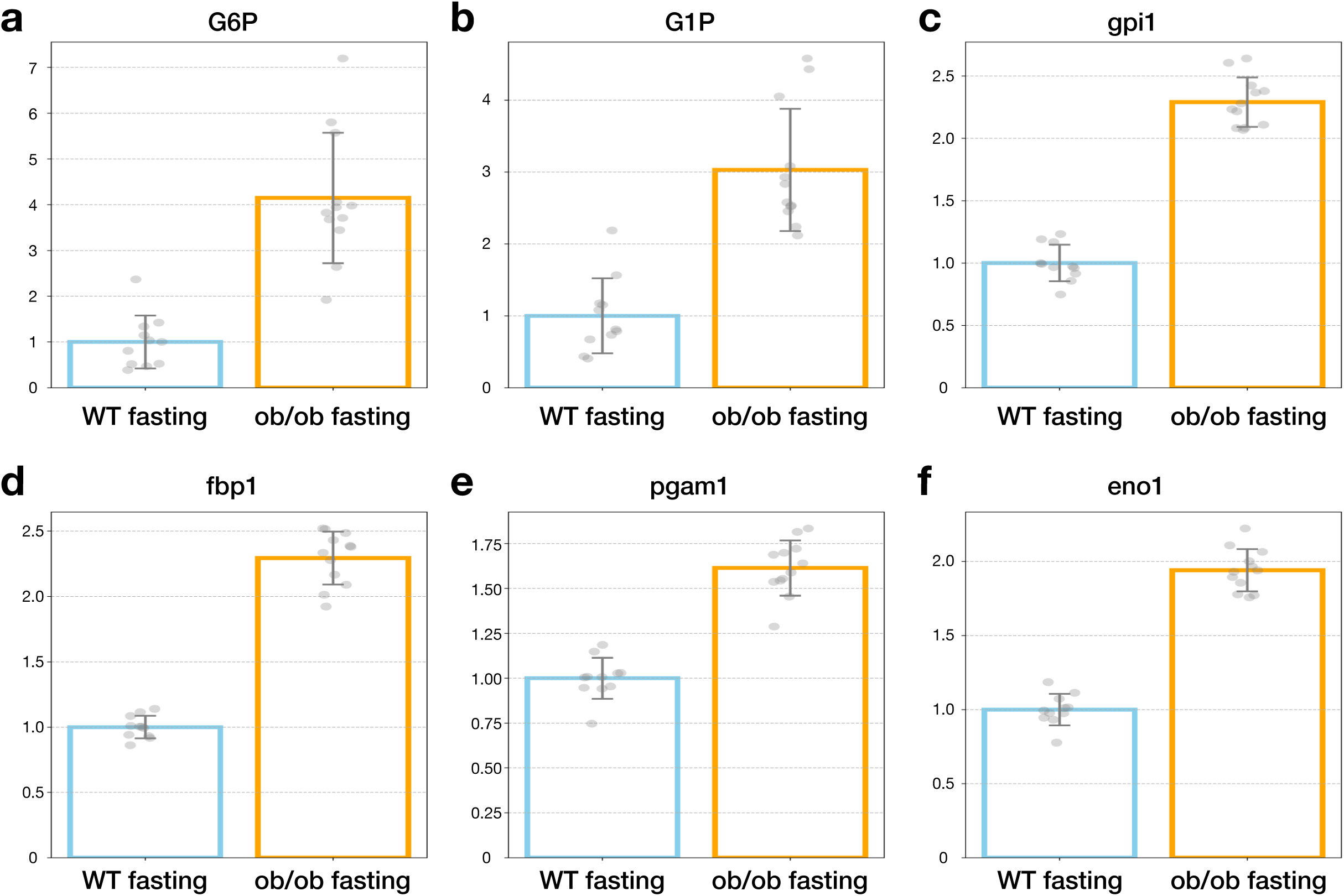
Relative expression of glycogenic enzymes and metabolites in mouse liver dataset. Relative expression of enzymes and metabolites, normalized to the mean value of the WT fasting group (set to 1). Each point represents an individual sample value (n=12 per condition), and the bars represent the mean value for each experimental condition. Error bars represent ±SD. (a) Glucose-6-phosphate (G6P), (b) Glucose-1-phosphate (G1P), (c) Glucose phosphate isomerase 1 (gpi1), (d) Fructose-1,6-bisphosphatase 1 (fbp1), (e) Phosphoglycerate mutase 1 (pgam1), (f) Enolase 1 (eno1)

